# Linking nutrient stoichiometry to Zika virus transmission in a mosquito

**DOI:** 10.1101/645911

**Authors:** Andrew S. Paige, Shawna K. Bellamy, Barry W. Alto, Catherine L. Dean, Donald A. Yee

## Abstract

Food quality and quantity serve as the basis for cycling of key chemical elements in trophic interactions, yet the role of nutrient stoichiometry in shaping host-parasite interactions is under appreciated. Most of the emergent mosquito-borne viruses affecting human health are transmitted by mosquitoes that inhabit container systems during their immature stages, where allochthonous input of detritus serves as the basal nutrients. Quantity and type of detritus (animal and plant) were manipulated in microcosms containing newly hatched *Aedes aegypti* mosquito larvae. Adult mosquitoes derived from these microcosms were allowed to ingest Zika virus infected blood and then tested for disseminated infection, transmission, and total nutrients (percent carbon, percent nitrogen, ratio of carbon to nitrogen). Treatments lacking high quality animal (insect) detritus significantly delayed development. Survivorship to adulthood was closely associated with the amount of insect detritus present. Insect detritus was positively correlated with percent nitrogen, which affected Zika virus infection. Disseminated infection and transmission decreased with increasing insect detritus and percent nitrogen. We provide the first definitive evidence linking nutrient stoichiometry to arbovirus infection and transmission in a mosquito using a model system of invasive *Ae. aegypti* and emergent Zika virus.

## INTRODUCTION

The environment experienced during development often greatly shapes adult phenotypes. For animals with complex life cycles, ecological factors during the juvenile stage can influence the adult stages via carry-over effects, including in fish (Green and McCormick 2005, McCormick and Gagliano 2008), mosquitoes (Alto et al. 2005), and other insects (Kingslover et al. 2011). In insects, and mosquitoes in particular, different temperatures, habitats, and food environments can affect different life history states, and thus adults may result with differential contributions to overall lifetime fitness (Kingslover et al. 2011). For mosquitoes, this means that larvae exposed to different stressors may have different adult life history traits including mass, development time, survivorship to adulthood, and population growth, as well as and resilience to future stress including pathogen challenge (Nasci 1991, Alto et al. 2012, Da-Silva Araújo et al. 2012). These life history traits in addition to vector competence, defined as the susceptibility to pathogen infection and transmission potential, are some of the many factors that influence vectorial capacity. Vectorial capacity is an index of risk of pathogen transmission (Alto et al. 2008, Bara et al. 2014), and if defined as the number of infectious bites a host receives per day.

Nutrition is an early developmental factor known to modulate life history traits. It is an important factor regarding host-pathogen interactions; as host nutrition influences multiple measures of adult performance, immune response, and the resources necessary for pathogen replication (Lee et al. 2006, 2008; Telang et al. 2012, Yee et al. 2015). Larval nutritional stress can reduce the immune response of mosquitoes allowing for increased pathogen infection (Sindbis virus; Muturi et al. 2011) or reduce resources for pathogen development (malaria; Vantaux et al. 2016ab). Further, nutritionally stressed larvae result in adults with reduced mass. As larger bodied mosquitoes imbibe greater volumes of blood than their smaller counterparts, one may predict that large mosquitoes would ingest higher doses of pathogen in the infected blood and may have higher infection rates, as arboviral infection is often dose dependent (Takken et al. 1998, McCann et al. 2009).

Past studies have explored the role of nutrition in the larval environment primarily in terms of stress induced by varying quantities of laboratory or plant detritus as a substrate for microbial growth; these microbes provide the direct source of larval nutrition. Recent studies utilizing stable isotope analysis have shown that additions of animal detritus increase nitrogen availability. In particular, increases in animal detritus have positive effects on larval growth and adult phenotypes such as decreasing development time, larger mean mass, greater survivorship to adulthood, and higher estimated population growth (Yee and Juliano 2006, Murrell and Juliano 2008, Winters and Yee 2012, Yee et al. 2015). Within natural and artificial aquatic container systems such as treeholes and tires (communities dominated by immature stages of mosquitoes), primary production is nearly absent. Most of the incoming energy originates from allochthonous inputs of detritus, mainly in the form of senescent plant material (primarily leaves) and terrestrial invertebrate carcasses (Carpenter 1983, Kling et al. 2007). Invertebrate carcasses, which make up the bulk of animal detritus, have greater available nitrogen stores and a faster rate of decay than plant material, allowing for more rapid release of nutrients into the system (Yee and Juliano 2006). This suggests that animal detritus scarcity could have important effects on vector competence as a limiting factor of growth, impacting a variety of adult traits, including immunity.

At the molecular level, immunity has several components including the production of cells that actively destroy pathogens. For example, within this pathway, nitric oxide (NO) plays a critical role in innate immunity in both vertebrates (Wink et al. 2011) and insects like mosquitoes (Hillyer & Estéves-Lao 2010). Free radicals like NO are very unstable and react quickly with other molecules to acquire a stable electron configuration (Clements 2012). As an important component in immunity, it seems possible that nitrogen limitation could affect immunity against pathogens. Specifically, reactive nitrogen is linked to mosquito immunity as nitrate and hydrogen peroxide are used to synthesize NO in the mosquito midgut (Hillyer 2010).

In this study, we aimed to investigate the influence of various ratios of animal:plant detritus on infection and transmission of Zika virus in *Aedes aegypti*. We test the hypothesis that nutrient limitation (specifically Nitrogen limitation) during the larval stages will be associated with higher infection and transmission potential of *Ae. aegypti* for Zika virus. We measure total nutrients in terms of percent carbon (%C), percent nitrogen (%N), and ratio of carbon to nitrogen (C:N) in both the mosquitoes and basal resources (detritus). Inclusion of measurements of both detritus and mosquitoes allows us to provide a link between basal nutrition and the adult phenotype, in terms of nutrient stoichiometry. Although %C and %N are correlated to C:N, the latter value is reflective of the stoichiometry of the animal, in essence showing how they may balance the two elements in their body across detrital environments. Although carbon and nitrogen are contained within C:N, individually they cannot enlighten us about how they act in tandem. Although we focus on *Ae. aegypti* and the Zika model system, this study has general application in addressing a gap in our understanding of how mosquito larval nutrition relates to adult nutrient stoichiometry and interactions with pathogens ingested from infected vertebrate blood.

## MATERIALS AND METHODS

### Mosquitoes and Detritus Treatments

*Aedes aegypti* mosquitoes used in these studies were collected as larvae from containers in Key West, FL. Larvae were reared in approximately 1.0 L of tap water in plastic trays (25 x 30 x 5 cm) with 0.40 g larval food comprised of equal amounts of liver powder and brewer’s yeast at hatching and supplemented with the same amount of food 3-4 d later. Pupae were transferred to water-filled cups in 0.3 m^3^ screened cages for emergence to adulthood. Adults were provided with 10% sucrose solution from cotton wicks and weekly blood meals from live chickens (IACUC protocol 201507682). Females laid eggs on damp paper towels in cups with water held in the cages. All life stages were maintained with a light:dark photoregime of 12:12 h at 28°C. The F_17_ generation of parental *Ae*. *aegypti* were used in these experiments.

Larval rearing treatments consisted of ten groups that varied in the relative ratio and amount of animal (freeze-dried crickets, *Gryllodes sigillatus*, Fluker Farms, Port Allen, LA, USA) to plant (senescent red maple leaves, *Acer rubrum,* collected at the Lake Thoreau Center, Hattiesburg, MS, USA 31°19’37.63”N, 89°17’25.22”W) detrital sources. Plant and animal detritus were dried for 48 hrs at 45 °C prior to use. Each detritus treatment was performed in triplicate for a total of 30 experimental units (hereafter, containers). Detritus types were expressed in relative terms (1 unit of detritus equals 0.15 g) of animal:plant as follows: 1:0, 2:0, 4:0, 0:5, 0:10, 1:5, 1:10, 2:5, 2:10, 4:10. These detritus treatments allow for a range of nitrogen and carbon values in adults and generally were based off past studies examining how different detrital environments affect mosquito performance and stoichiometry (e.g., Winters and Yee 2012, Yee et al. 2015). To permit microbial growth for mosquito larvae to feed on, detritus was soaked for 5 d before introduction of larvae in 2.0 L plastic buckets (height, 19.05 cm; top and bottom diameters, 19.30 cm and 16.31 cm, respectively) containing 2000 mL tap water and 1000 µL of tire water inoculum. Tire water inoculum was obtained from several tires occupied by mosquitoes and maintained on the UF-FMEL campus in Vero Beach, FL. The inoculum provided a source of microorganisms, acquired from a semi-natural setting, as food for larvae. Treatment containers were maintained with a light:dark photoregime of 12:12 h at 28°C.

Eggs were hatched at room temperature for 60 min in a deoxygenated water in a 250 mL Erlenmeyer flask attached to a vacuum to induce synchronous hatching with 0.20 g/L of larval food (Kauffman et al., 2017). Larvae were transferred to 5 L of tap water in 5 L plastic trays with an additional 0.20 g/L food. The following day, the first instar larvae were rinsed with tap water to remove larval food and 160 larvae were placed in each treatment container. The initial larval density (0.08 larvae/mL) is within the range of densities observed in field conditions in Florida among tires occupied by *Ae. aegypti* and competitor *Ae. albopictus* (Alto et al. 2005). Treatment containers were maintained in a walk-in environmental chamber at the UF-FMEL with a light:dark cycle of 12:12 h set at 28±1°C. Containers were checked every day and rearranged haphazardly within the environmental chamber. When present, pupae were transferred from treatment containers to polystyrene *Drosophila* culture vials with 2 to 5 mL of tap water (up to 5 pupae per tube) according to treatment conditions and sealed with a cotton plug. The date and sex of newly emerged adults from each replicate were recorded. Both males and females were housed together by treatment, replicate, and emergence date in paperboard cages with mesh screening (height by diameter: 10 cm x 10 cm) and rearranged haphazardly each day in the environmental chamber. For logistical reasons, females were added into cages over a period of three days because it would have been impractical to blood feed large numbers of cages. Adults were provided with 10% sucrose solution on cotton pads. Females were 9 to 15 d old at the start of trials in which mosquitoes were allowed to ingest Zika virus infected blood.

### Infection Study

Females were fed defibrinated bovine blood (Hemostat Laboratories, Dixon, CA) containing freshly propagated Zika virus. To encourage blood feeding, mosquitoes were deprived of sucrose, but not water, 24 h before blood-feeding trials. Infection experiments were performed in a biosafety level-3 laboratory at the UF-FMEL. Isolates of Asian lineage of Zika (strain PRVABC59, GenBank KU501215.1) from Puerto Rico were prepared in African green monkey (Vero) cells and used in the infection study. Monolayers of Vero cells were inoculated with 500 µL of diluted stock virus (multiplicity of infection, 0.1) and incubated at 1 h at 37 °C and 5% CO_2_ atmosphere, after which 24 mL media (M199 medium supplemented with 10% fetal bovine serum, penicillin/streptomycin and mycostatin) were added to each flask and incubated for 4 d. Freshly harvested media from infected cell cultures were combined with defibrinated bovine blood and adenosine triphosphate (0.005 M) and presented to mosquitoes using an artificial feeding system (Hemotek, Lancashire, UK) with hog casing membranes for 1 hr feeding trials. Carbon dioxide from the sublimation of dry ice was used to stimulate blood feeding three times every 20 min. Samples of infected blood were taken at the time of feedings and stored at −80 °C for later determination of virus titer. Mosquitoes were fed 6.5 - 7.5 log_10_ plaque forming units (pfu)/mL of Zika.

Following blood feeding trials, fully engorged mosquitoes were sorted using light microscopy and held in cages, maintained at 12:12 hour L:D photoperiod and at 28 °C. Partially fed (average of 2%) and unfed females (average of 9%) were discarded. Mosquitoes were provided with an oviposition substrate and 10% sucrose solution on cotton pads.

### Zika virus Disseminated Infection, and Transmission Potential

Mosquito tissues were tested for Zika infection 15 d after ingestion of infected blood. Mosquito legs and saliva were tested for Zika RNA as indicators of Zika disseminated infection (Turell et al. 1984) and transmission potential, respectively (i.e., the presence of viral RNA in saliva is a proxy for transmission). Partitioning mosquito tissues for testing allowed us to determine treatment-induced changes in barriers to infection (e.g., midgut escape barrier and salivary gland barriers). Mosquitoes were cold anesthetized (4 °C), and the legs were removed using light microscopy. Legs were placed in 1 mL of incomplete media (M199) and stored at −80 °C until testing. Using forceps, one wing was damaged to immobilize the mosquito and the proboscis was inserted into a capillary tube for a 1-h collection of saliva in type B immersion oil using methods described by Alto et al. (2014). Following collection of saliva, mosquito bodies were stored at −80 °C until nutrient analysis testing. Saliva and oil were expelled into 300 μL of media (M199) and stored at −80 °C until testing. Each treatment replicate yielded multiple mosquitoes and so infection measures are reported as percent infection per replicate.

### RNA Extraction and Reverse Transcriptase Quantitative Polymerase Chain Reaction (RT-qPCR)

Legs and bodies were homogenized using a TissueLyser (Qiagen, Valencia, CA) in 1000 μL media after which a 140 μL sample of homogenate was clarified by centrifugation and used for RNA isolation with the QIAamp viral RNA mini kit (Qiagen, Valencia, CA) following the manufacturer’s protocol. Saliva samples were processed similarly, but with no homogenization. Viral RNA was eluted in 50 μL buffer and quantitative RT-PCR was used to determine the presence and quantity of viral RNA using the Superscript III One-Step qRT-PCR with Platinum® Taq kit (Invitrogen, Carlsbad, CA) with the C1000 Touch Thermal Cycler, CFX96 Real-Time System (Bio-Rad Laboratories, Hercules, CA). The mastermix used 2.2µL molecular grade water, 1µL forward primer, 1µL reverse primer, 10µL 2X reaction mix, 0.4µL ZIKV probe, 0.4µL Taq polymerase, and 5µL of mRNA template. Primers and probe sets synthesized by IDT (Integrated DNA Technologies, Coralville, IA) had the following sequences: Forward Primer, 5′-CTTCTTATCCACAGCCGTCTC-3′ Reverse Primer, 5′-CCAGGCTTCAACGTCGTTAT-3′ Probe, 5′-/56-FAM/AGAAGGAGACGAGATGCGGTACAGG/3BHQ_1/-3′

The program for qRT-PCR consisted of a 30-min step at 50°C linked to a 40-cycle PCR (94°C for 12 s and 58°C for 60 s). A standard curve was used to quantify viral load (titer) of Zika in mosquito tissues by comparing cDNA synthesis to a range of serial dilutions of Zika in parallel with plaque assays of the same dilution of virus, expressed as pfu equivalents/mL (Bustin 2000).

### Carbon and Nitrogen Analysis

Mosquito species can vary in nutrient content, both based on larval diet and inherent differences among species (e.g., Yee et al. 2015). Carbon is the principle building block of life, and can vary with across diet. In addition, as container systems for developing *Aedes albopictus*, like tree holes, are nitrogen limited (Carpenter 1983) we focused on percent body nitrogen as well. For nutrient analysis, mosquitoes and detritus were prepared by drying in an oven for at least 48 hrs at 50°C. Each weighed sample (mosquito and detritus) was encapsulated in 5 x 9 mm pressed tin capsules (Costech Analytical, Valencia, CA, USA) before analysis. Mosquito body samples and detritus samples were analyzed for total nutrients (%C, %N, C:N) using a ECS 4010 Elemental Combustion System (Costech Analytical Technologies, California). Although %C and %N are correlated to C:N, the latter value is reflective of the stoichiometry of the animal, in essence showing how they may balance the two elements in their body across detrital environments. Although carbon and nitrogen are contained within C:N, individually they cannot enlighten us about how they act in tandem.

### Estimated Finite Rate of Increase

In many cases, life history traits correlate with per capita rate of change. An estimate of the per capita rate of change is feasible in experiments where populations are established as cohorts and indirect measures of survivorship, fecundity and generation time are available (Livdahl and Sugihara 1984, Juliano 1998). An estimate of the finite rate of increase (λ′) was calculated for each replicate container by initially calculating the estimated instantaneous rate of increase (r’, Livdahl and Sugihara 1984):

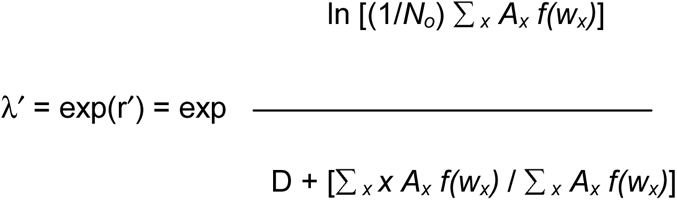

where *N_0_* is the initial number of females in the cohort (assumed to be 50%); *A_x_* is the number of females emerging to adulthood on day x; D is the time from emergence to reproduction taken as 12 d for *Ae. aegypti* (Grill and Juliano 1996); *f* (*w_x_*) is a function based on the relationship between size and fecundity in female mosquitoes. For *Ae. aegypti f* (*w_x_*) = 2.5 *w_x_* – 8.616 (Briegel 1990).

### Statistical Analysis

Analysis of variance (ANOVA) was used to test for larval treatment effects on male development time, survivorship to adulthood, and the estimate of the finite rate of increase (λ′). Multivariate analysis of variance (MANOVA) was used to test for treatment effects on adult female development time and mass. Standardized canonical coefficients were used to describe the relative contribution and relationship of the response variables to the multivariate treatment effect. Differences in response variables among treatment groups were identified using the Tukey-Kramer HSD post-hoc test for multiple comparisons. Stepwise multiple regression analysis was used to relate infection measures (disseminated infection, saliva infection) to detrital conditions (%N, amount of animal detritus) and %C, %N, C:N signatures in adult females (pooled across treatments). All statistical analyses were performed using SAS software (2004). A randomization ANOVA was used to analyze treatment effects on λ’ due to no transformations allowing us to meet assumptions of normality and heteroscedasticity.

## RESULTS

### Mosquito Life History

Multivariate analysis of variance showed significant effects of treatment on female development time and mass (Pillai’s trace _18,34_ = 1.72). Standardized canonical coefficients showed that development time contributed almost twice as much as mass to the significant treatment effect (SCCs, development time = −2.50, mass = 1.42). Females with the longest development times were associated with reduced mass (Fig. 1).

**Figure 1.**
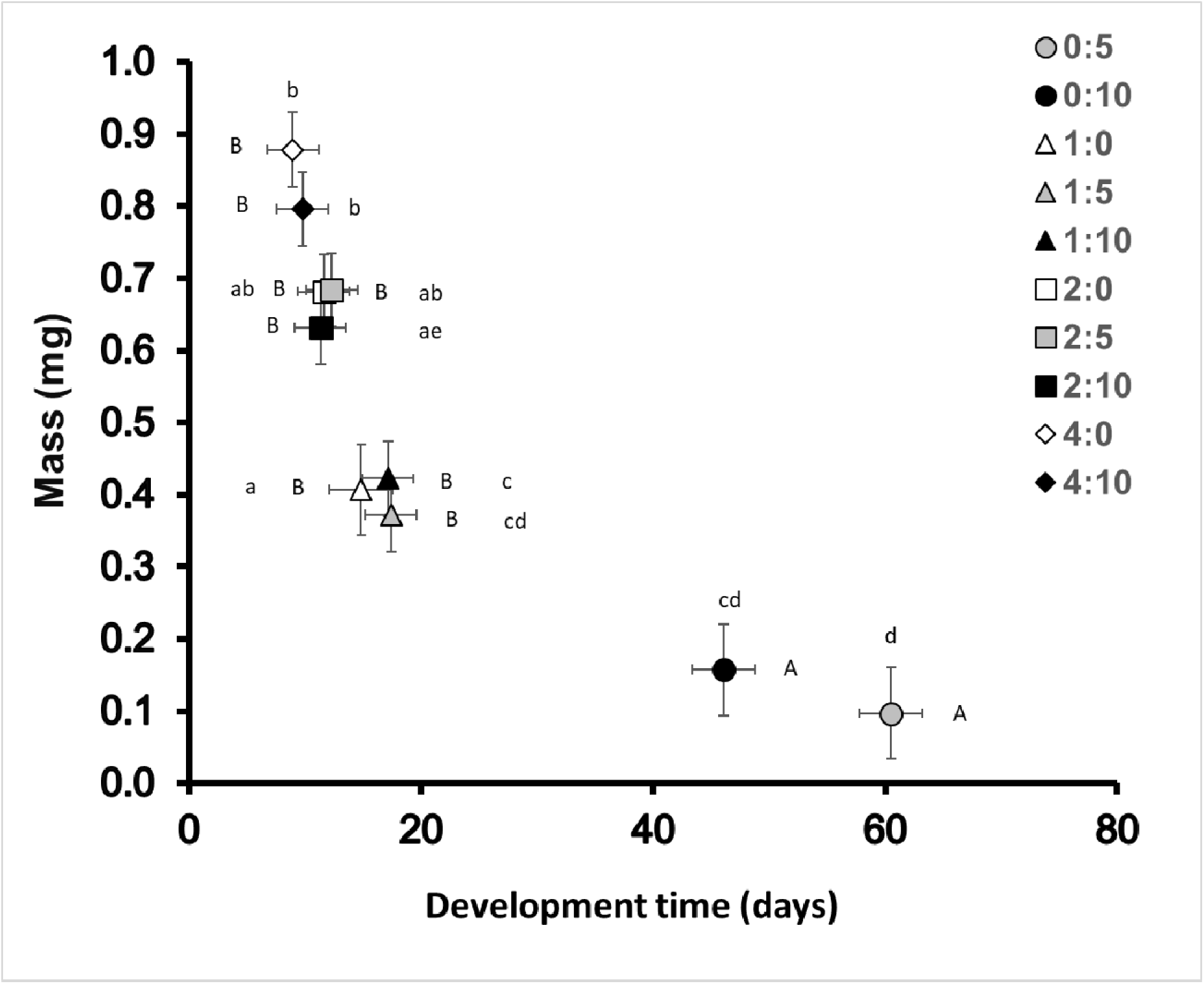
MANOVA of bivariate least square means of female mass and development time across different nutrient environments as represented by different ratios of animal (crickets) and leaf (red maple) detritus. Means that do not share same letters are significantly different, and bars indicate standard error of the mean. Lowercase letters are for mass and uppercase letters are for development time.

Treatment levels lacking animal detritus displayed significantly delayed development time for both female and male mosquitoes (F_8,25_ = 45.42, P < 0.001). Plant only treatments showed delayed development compared to treatment levels with animal detritus (with or without plant) (Figs. 1 and 2). No significant differences were found when treatment levels included at least one unit of animal detritus. No male survivors were observed in the treatment 0:5.

**Figure 2.**
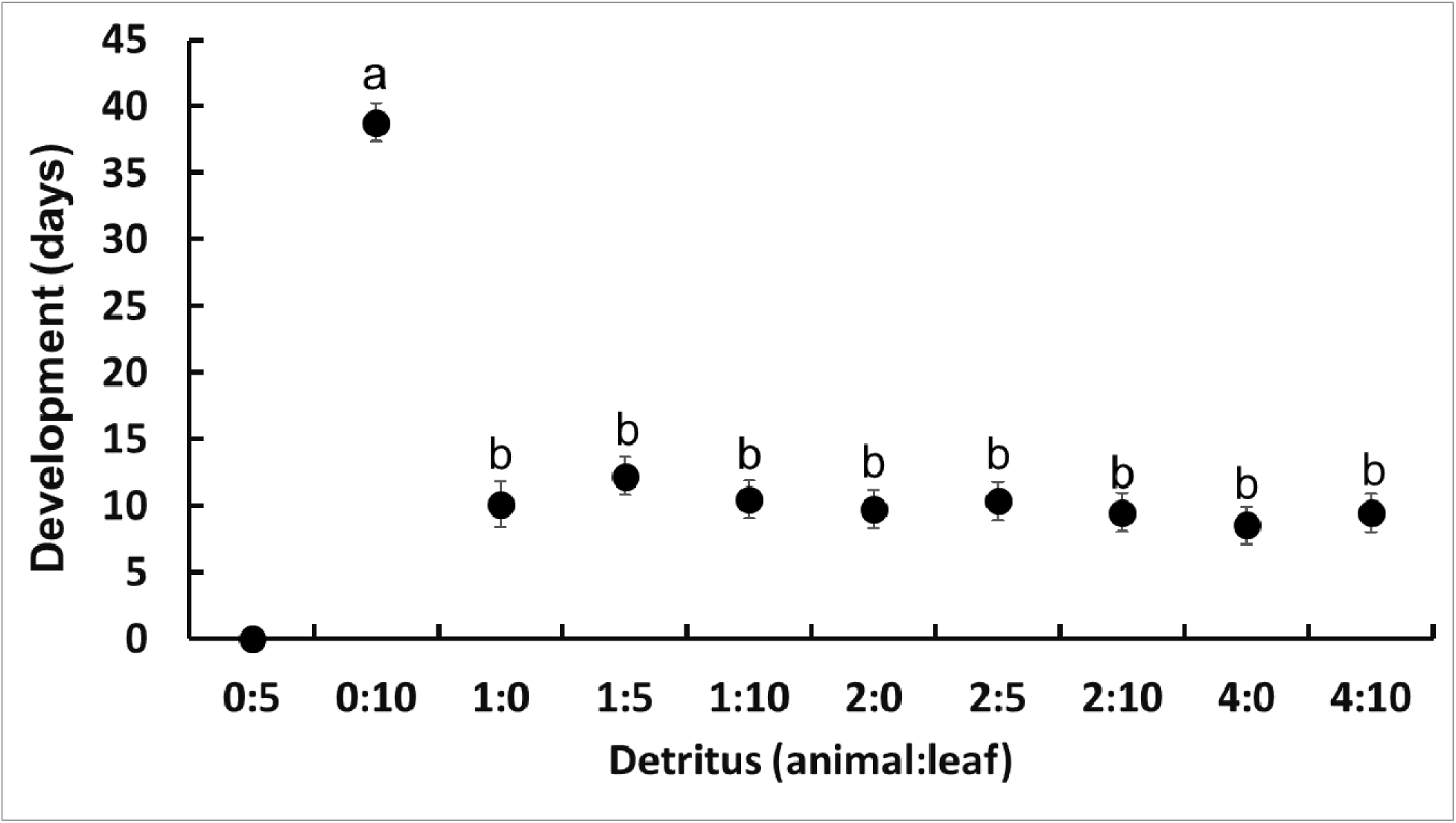
ANOVA of least square means of male development time across different environments as represented by different ratios of animal (crickets) and leaf (red maple) detritus. Means that do not share same letters are significantly different, and bars indicate standard error of the mean.

There was a significant effect of treatment on mosquito weights. Mosquitoes were the heaviest in treatment levels with at least two units of animal detritus, regardless of how much plant detritus was present (Fig. 1). Mosquitoes were intermediate in weight with one unit of animal detritus, regardless of the amount of leaf detritus was preset (Fig. 1). The lightest mosquitoes were produced in habitats with only leaf detritus present (Fig. 1).

Survivorship to adulthood was closely associated with the amount of animal detritus present (F_9,27_ = 10.53, P<0.001). Increases in basal resources, especially inclusion of animal detritus, yielded higher survivorship compared to plant detritus only situations (Fig. 3). The highest survivorship was observed in treatments containing 2:5 and 2:10 units of animal:plant detritus. An intermediate to high survivorship was seen in treatments containing 1:10, 2:0, 4:0, and 4:10 units of animal:plant detritus, and intermediate to low survivorship in treatments containing 1:0 and 1:5 units of animal:plant detritus. The lowest survivorship was seen in treatments lacking animal detritus (Fig. 3).

**Figure 3.**
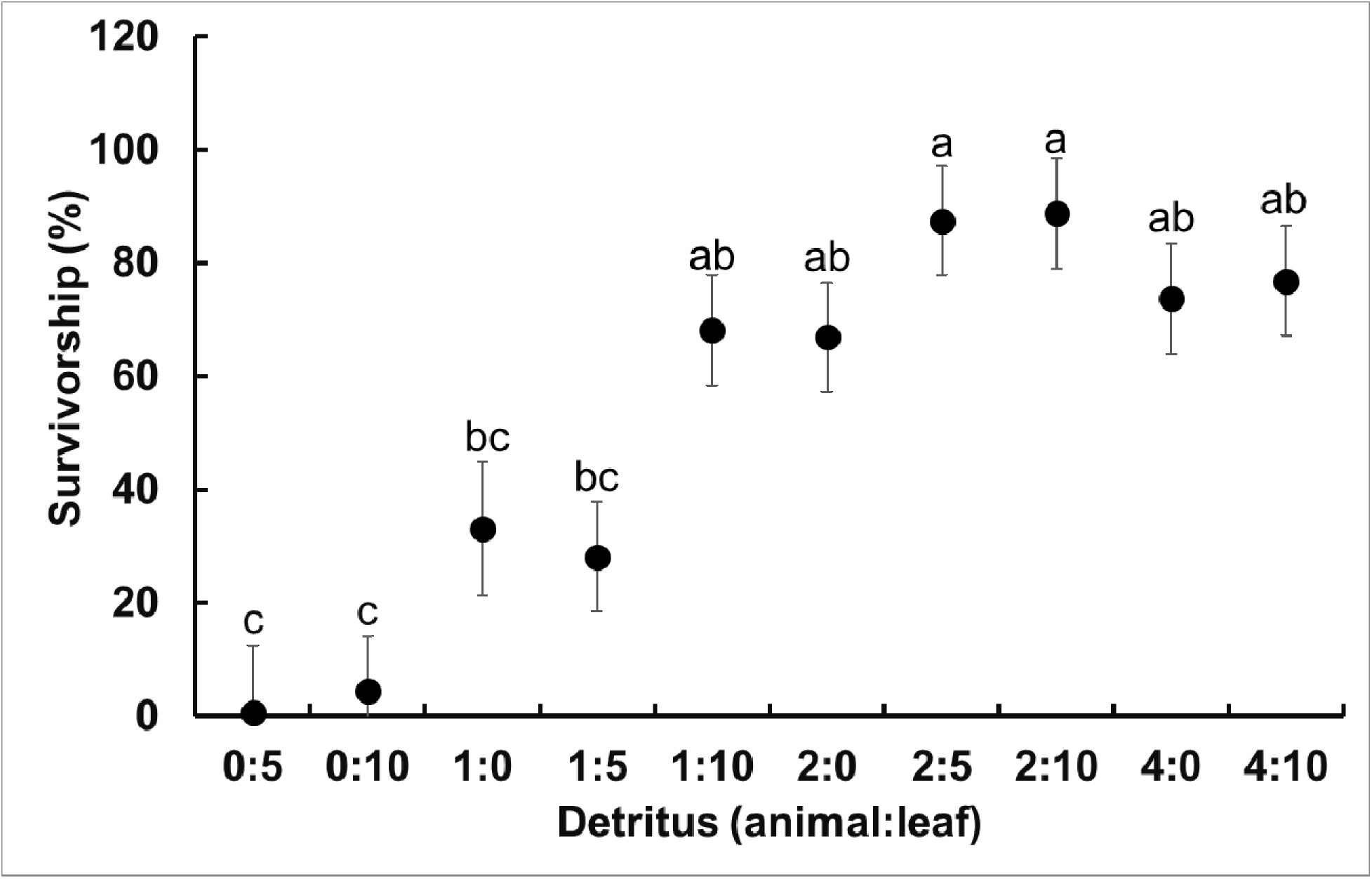
ANOVA of least square means of percent survivorship to adulthood (male+female, expressed as percent total of initial cohort of larvae added to containers) across different nutrient environments as represented by different ratios of animal (crickets) and leaf (red maple) detritus. Means that do not share same letters are significantly different, and bars indicate standard error of the mean.

A randomization ANOVA showed marginally non-significant treatment effects on λ′ (F_9,19_ = 2.28, P = 0.062, Fig. 4). In general, λ′ values were significant higher in combinations of animal and leaf detritus compared to leaf-only treatment levels. In most cases populations were estimated to be growing (λ′ > 1).

**Figure 4.**
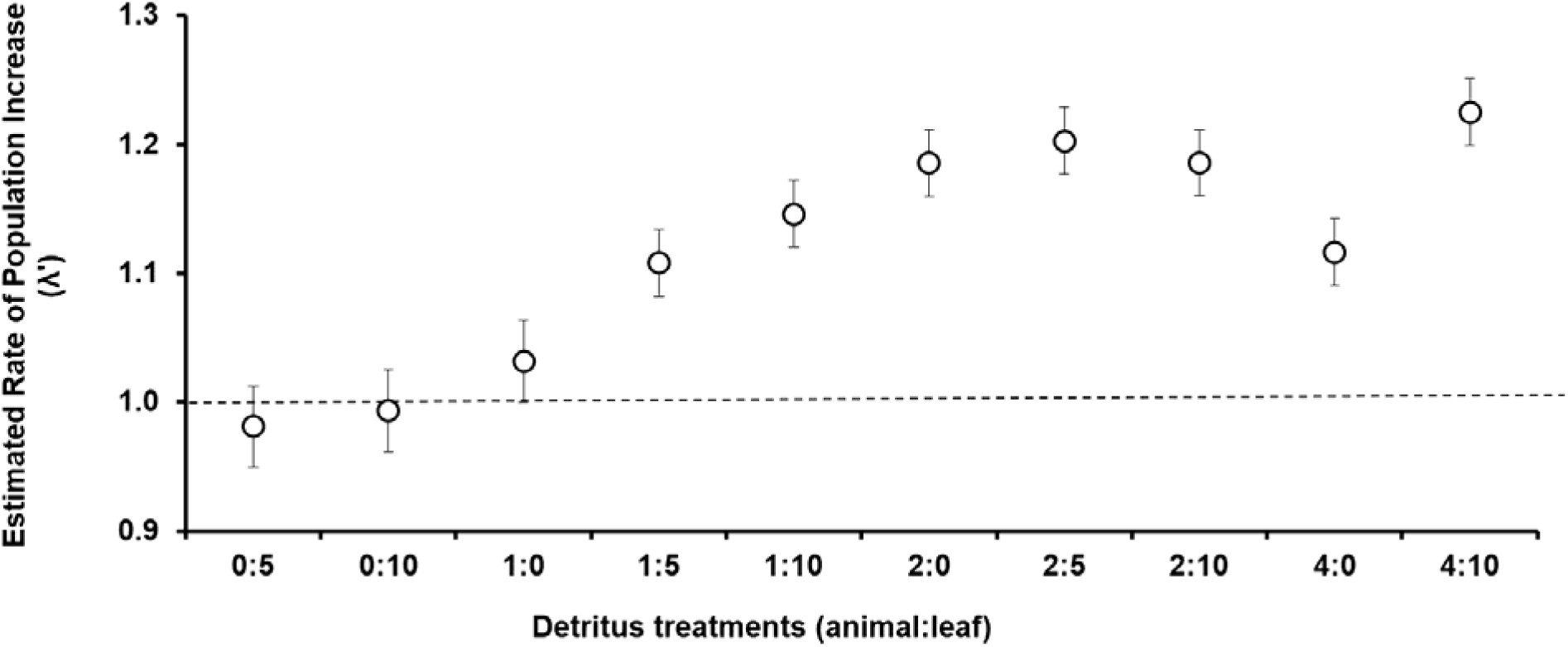
Values of the estimate of the finite rate of increase (λ′) for *Aedes aegypti* females across animal and leaf environments. Nutrient environments are represented by different ratios of animal (crickets) and leaf (red maple) detritus. Values that share a letter are not significantly different at P > 0.05. The dashed line at λ′ represents populations for which growth is estimated to be zero.

### Zika Virus Disseminated Infection and Transmission

Disseminated infection (stepwise regression: animal, F_1,20_ = 65.44, P < 0.001, R^2^=0.75; animal+leaf, F_2,20_ = 4.54, P = 0.046, R^2^= 0.05; %N, F_1,24_ = 9.23, P = 0.006) and transmission potential (stepwise regression, F_1,22_ = 20.30, P < 0.001) decreased with increasing animal detritus and %N (Figs. 5, 6, and 7). Disseminated infection was highest with treatment levels containing only one unit of animal detritus, intermediate in treatment levels containing two units of animal detritus, and low in treatment levels containing four units of animal detritus. Overall, adult females showed an average of 4.66 ± 0.09% nitrogen, 54.39 ± 0.54% carbon, and a 12.20 ± 0.26% C:N ratio across all detritus ratios.

**Figure 5.**
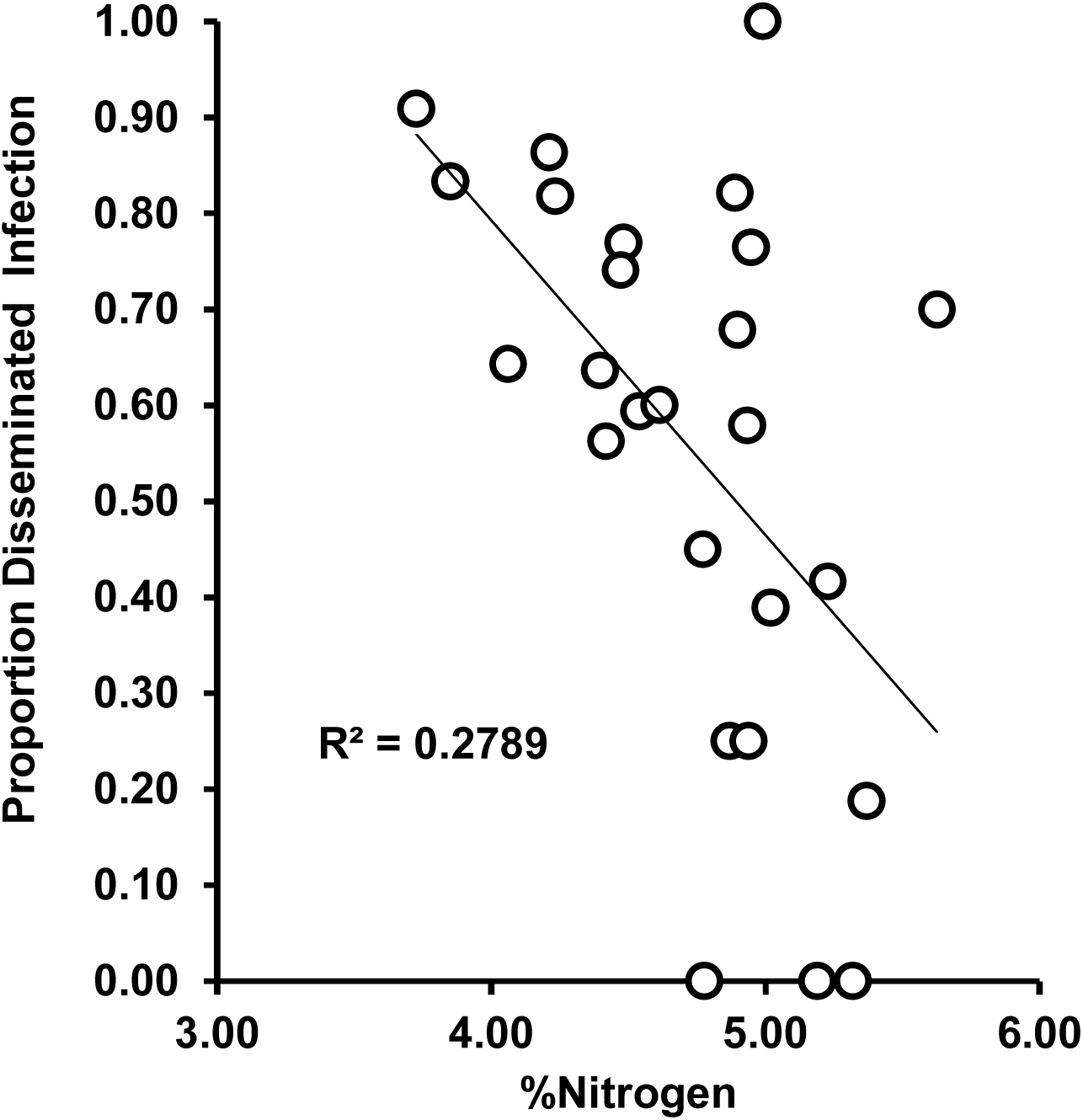
Stepwise multiple regression (%C, %N, C:N) on the proportion of positive mosquitoes in each treatment. Each point represents a replicate for each treatment (%N, F_1,24_ = 9.23, P = 0.006).

**Figure 6.**
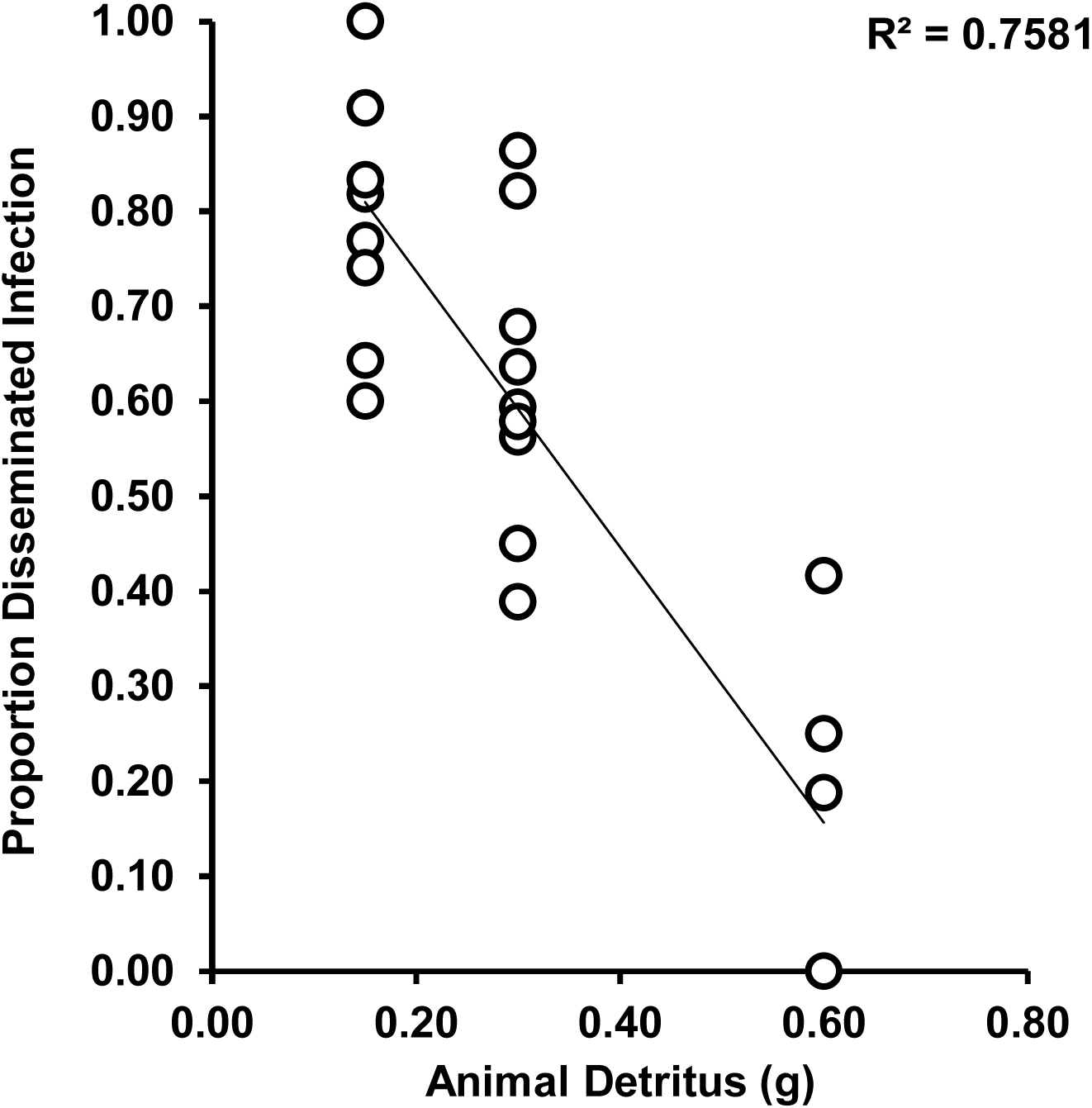
Stepwise multiple regression (%C, %N, C:N) on the proportion of positive mosquitoes in each treatment. Each point represents a replicate for each treatment (%N, F_1,24_ = 9.23, P = 0.006).

**Figure 7.**
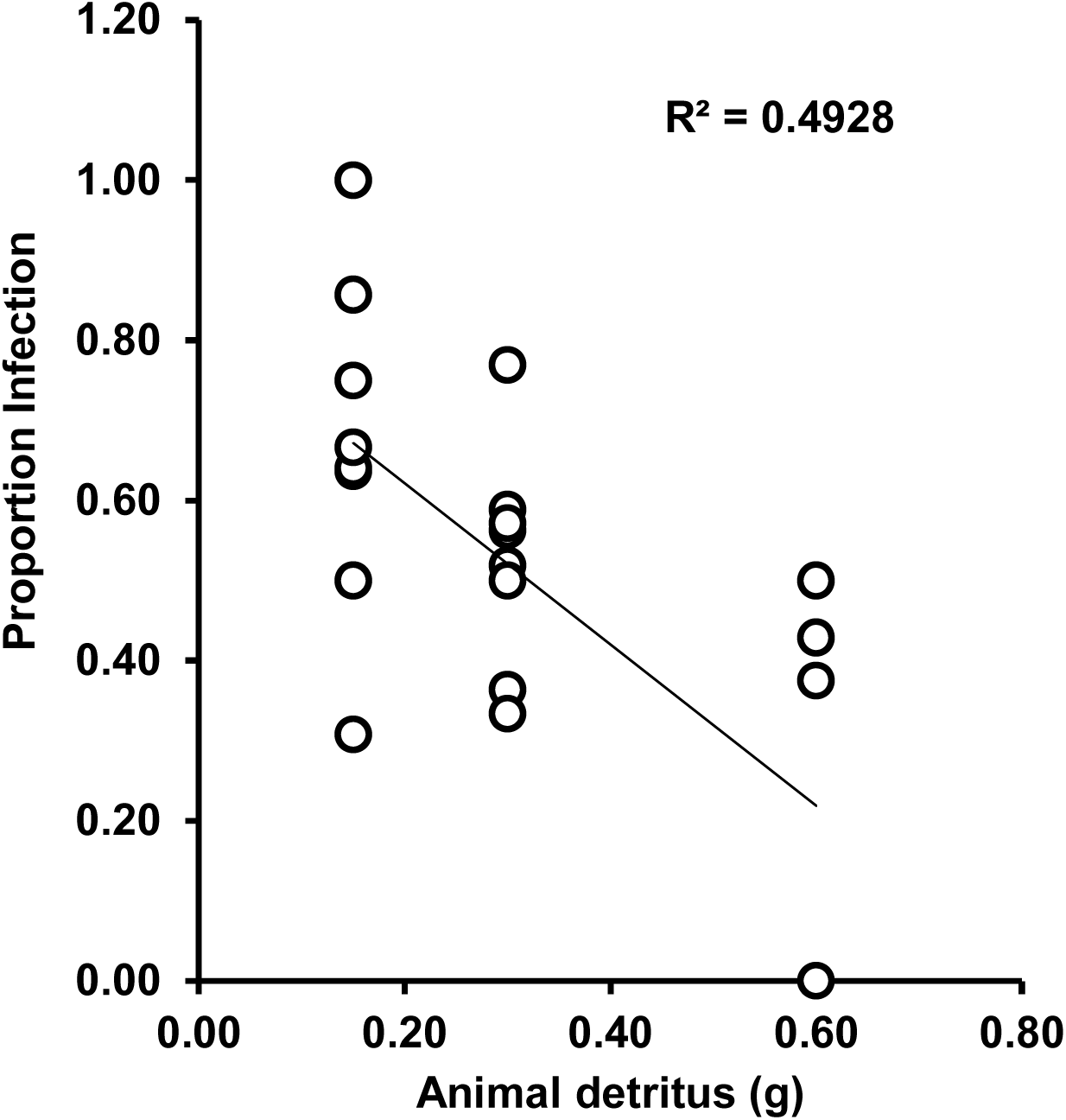
Stepwise multiple regression (animal detritus and leaf detritus (g)) on the proportion of mosquitoes with positive saliva infection in each treatment. Each point represents a replicate for each treatment (Animal, F_1,22_ = 20.30, P < 0.001, R^2^ = 0.4617; Leaf, not significant).

## DISCUSSION

We found support for our hypothesis that variation in animal and leaf detritus would alter Zika virus infection and transmission by *Ae. aegypti*. The infection component of our study revealed that quantity and quality of nutrition, and the associated changes in nutrient stoichiometry, altered disseminated infection and transmission potential of Zika virus. Particularly, animal detritus was positively correlated with %N, which affected Zika virus infection. Disseminated infection and transmission decreased with increasing animal detritus and %N. Thus, we provide the first definitive evidence linking nutrient stoichiometry to arbovirus infection and transmission in a mosquito, using a model system of *Ae. aegypti* and Zika virus. Future studies should consider using lower generation of mosquitoes (e.g., F1 generation from field-collected parents) which are likely to be more representative of populations in the wild. The observed resource quality mechanism mediating interactions between *Ae. aegypti* and Zika may apply to other arboviruses and mosquito species. For instance, stoichiometric composition was similar for both *Ae. aegypti Ae. albopictus* across different diet environments (Yee et al. 2015). These two species are often implicated in the same transmission cycles (e.g., chikungunya, Zika, dengue; Gubler 1998, Coffey et al. 2014, Boyer et al. 2018), so further work will be needed to determine if our findings of the relationships between stoichiometry and viral infection are applicable to *Ae. albopictus*. Although the mechanism for the observed results in the current study is unclear, it may relate to increased immune activity and reduced pathogen propagation, as observed in other systems (Cotter et al. 2011, Cornet et al. 2014, Brunner et al. 2014, Howick and Lazzaro 2014).

We were interested in producing females who would exhibit a range of nitrogen and carbon values and thus we used different combinations of high-quality animal and low-quality leaf detritus. Although we did produce a gradient (4.27 – 5.29 %N across diets), our diets yielded nitrogen values at the lower end of those produced elsewhere. For instance, in a laboratory experiment Yee et al. (2015) produced female *Ae. aegypti* with a range of 7.60 - 10.19 %N across diets using the same types but higher amounts of detritus per individual. Carbon levels were more similar, with our study producing adults with 51.38 – 58.78 %C whereas Yee et al. (2015) had a range of 45.75 - 55.45 %C. Thus, our diets produced females that were likely more stressed for limited nitrogen, although at present there are no published data from wild mosquitoes to know if the lower values produced in this study fall within the typical range for field collected adults. However, as our animals seemed more stressed for nitrogen and females in higher nitrogen containers had lower average disseminated infection, this does suggest that nitrogen does play a role in affecting vector-pathogen interactions; a mechanism which has not been explored elsewhere.

We observed that variation in the amount and relative ratio of animal to plant detritus altered individual life history trait responses of mosquitoes including development time to adulthood, mass (net growth), and survivorship to adulthood. Greater amounts of animal detritus and %N consistently shortened development time and resulted in heavier adults with higher survivorship. Thus, we were able to demonstrate that %N reflected, in presence of animal detritus, affected a variety of life history traits and rate of female infection. These observations are consistent with a study that demonstrated reduced development time and increased adult mass for *Ae. aegypti* and *Ae. albopictus* but not *Culex quinquefasciatus* in animal versus leaf only environments (Yee et al. 2015). Nutrient analyses showed that *Aedes* tissues varied in their C:N ratio dependent on animal and leaf detritus ratios, whereas *Cx. quinquefasciatus* showed a less plastic response in C:N ratio (Yee et al. 2015). This suggests that nutrient content, and not type of detritus, influenced life history traits.

In the current study, an estimate of the finite rate of increase (λ′) showed trends for different responses to the quantity and quality of nutrition. Specifically, the presence of animal detritus, either alone or in combination with leaf detritus, increased population growth relative to treatments with only leaf detritus present. However, this effect was only marginally non-significant. We hypothesized that increased variance attributable to treatments that had no survivors (λ ′= 0) was, in part, responsible for lack of significance as the low nutrient treatments showed drastically delayed development time and reduced survivorship. To test this hypothesis, we re-ran the analysis omitting replicates with no survivors (i.e., where λ ′= 0). Results showed a significant treatment effect (F _9,17_ = 8.77, P < 0.001) in the anticipated direction, despite reductions between treatment means among treatments, thus providing support for the hypothesis that increased variance was a contributing factor to the marginally non-significant result. This trend should not be taken lightly, as nutrient pulses within microcosms such as tree holes or tires are common. Further, spatio-temporal pulses of nitrogen in the form of animal detritus may account for rapid flux in mosquito populations with varying competence and longevity affecting disease transmission dynamics (Kling et al. 2007, Yee 2008, Yee and Juliano 2012).

Frost et al. (2008) observed rich nutrition in *Daphnia magna* water fleas enhanced growth and reproduction of a bacterial parasite (*Pasteuria ramosa*) and Vantuax et al. (2016b) reported a lesser likelihood of infection in females exposed to a reduced quantity of laboratory diet in the larval stages. Discrepancies in these observations may be, in part, attributable to the notion that elemental nutrition may alter parasite and pathogen infection dynamics at different stages of the infection cycle (Alto et al. 2015, Borer et al. 2016). Further, living pathogens, such as malaria parasites or *P. ramosa*, must acquire nutrients from the host environment to grow and reproduce whereas viruses must hijack host replication machinery to replicate. Calorie restriction has been shown to either decrease or increase resistance to parasitism (reviewed in Cotter et al. 2011). The mechanism(s) may be, in part, related to the observation that different immune traits (e.g., phenoloxidase activity and lysozyme-like antibacterial activity) respond differently to nutrient uptake, as demonstrated in the Egyptian cotton leafworm (Cotter et al. 2011). At present, the mechanism for why *Ae. aegypti* females would be less susceptible to infection by Zika when nitrogen levels are greater is unclear.

However, given that container systems that produce adults are often limited by nitrogen (Carpenter 1983), this area of research could prove fruitful for linking fine-scale patterns of resource environments to human outbreaks of arbovirus induced disease. Although the present study was limited by the amount of tissue required to perform elemental analysis for C, N, and P, future investigations should include of the role of phosphorus and other essential elements to understand the role of limiting nutrients in infection.

Our study showed that infection traits map onto different regions of nutrient space as observed by other studies (Cotter et al. 2011). The effects of dietary intake on nutrient stoichiometry and subsequently on immunity in insects has only been investigated in a handful of disparate taxa (Lee et al. 2008). However, it is known that activation or maintenance of immunity often involves use of protein reserves (Lee et al. 2006), which is likely consistent with nitrogen availability. Thus, our results provide a starting point to investigate the wider role of nutrients, including nitrogen, in affecting mosquito-pathogen interactions of important human diseases.

Adult mosquitoes derived from nutrient rich environments containing insect detritus had greater mass and body size than adults from treatments with less detritus, especially those with little or no animal detritus. Although larger mosquitoes consume greater volumes of blood, therefore ingesting higher doses of Zika, we considered the possibility that they might have higher rates of infection. However, the pattern that we observed was the opposite of this prediction. Specifically, larger adults from nutrient rich larval environments had lower disseminated infection and saliva infection rates than smaller conspecifics. This observation suggests that differences in infection rates are not attributable to differences in volume of infected blood ingested. Rather, we speculate that the overall health of the mosquito determined by larval nutrient environments, may influence infection and progression of infection (advanced states of infection). Another possibility is that larger blood meals may provide a greater influx of nitrogen. Consequently, some blood meal resources for reproduction may trade off with energy reserves to fight an infection in order to live long enough to reproduce successfully which would likely be adaptive and favored by selection. Other studies investigating mosquito larval nutrition (amount of plant detritus or quantity of laboratory diet) and competition have observed similar impacts on adult life history traits or competence (LaCrosse virus, Grimstad and Haramis 1984; Sindbis virus, Alto et al. 2005; dengue virus, Alto et al. 2008ab). However, we are the first to quantify nitrogen limitation and demonstrate it’s role in arboviral infection and transmission potential. Invasive *Ae. albopictus* and native *Ae. triseriatus* container mosquitoes derived from nutrient rich larval environments were less likely to exhibit disseminated infection and/or to transmit dengue-2 virus (Zhang et al. 1993) and LaCrosse encephalitis virus (Grimstad and Haramis 1984, Grimstad and Walker 1991, Paulson and Hawley, 1991), respectively, than conspecifics from nutrient-deprived larvae. Additionally, these nutrient effects carried-over to the next generation as demonstrated with maternal effects on offspring infection with LaCrosse virus. This may have important epidemiological consequences given that vertical transmission is a mechanism for persistence of LaCrosse in the environment (Patrician and DeFoliart 1985). Thus, we propose that larval diet, with specific reference to the nutrients it contains, is a mechanism that affects nutrient composition and allocation patterns in *Ae. aegypti*; it may be an important but overlooked component to understanding transmission potential of arboviruses across different resource environments.

Larval nutrition alters several phenotypic traits related to mosquito fitness that are relevant to their ability to transmit pathogens (Beldomenico and Begon 2010) such as longevity (Steinwascher 1982, Haramis 1985), host-seeking behavior (Nasci 1986, Klowden et al. 1988), biting persistence (Nasci 1991), blood-feeding success (Nasci 1986), and fecundity (Steinwascher 1982, Vantaux et al. 2016a). It is likely that enhanced infection associated with nutrient deprivation may also have consequences for these other life history traits, so mathematical models are needed to evaluate the net effect on risk of arbovirus transmission (Bara et al. 2014).

## ACKNOWLEDGMENTS

An isolate of Zika virus was kindly provided by the Centers for Disease Control and Prevention. We thank A. Carels, B. Eastmond, S. Ortiz, K. Wiggins, and R. Zimler for technical support with experiments. K. Kuehn provided technical support for nutrient analysis of mosquitoes.

